# Plasma metabolomics reveals lipid-predominant metabolic disruptions in virologically suppressed people with HIV

**DOI:** 10.64898/2026.08.19.745653

**Authors:** Christopher M. Basting, Candace Guerrero, Kevin Escandón, Jodi Anderson, Garritt Wieking, Erik Swanson, Ty Schroeder, Charlotte Hemmila, Ross T. Cromarty, Fernanda Torres-Ruiz, Maribel Soto-Nava, Lady Carvajal-Ruiz, Karla Krystel Ordaz-Candelario, Olivia Briceño, Nicholas Funderburg, Santiago Avila-Rios, Melanie L. Graham, Timothy W. Schacker, Gonzalo Salgado Montes de Oca, Nichole R. Klatt

## Abstract

People with HIV (PWH) on antiretroviral therapy (ART) experience excess morbidity and mortality from comorbidities including cardiovascular and metabolic disease, yet the biological mechanisms underlying these outcomes in virally suppressed PWH (VS-PWH) remain incompletely understood. We applied high-dimensional targeted plasma metabolomics to quantify 254 small molecules and 750 lipids across 49 classes in viremic PWH (Vi-PWH), VS-PWH, and people without HIV (PWoH), analyzed alongside multiple T cell metrics and plasma cytokine concentrations. Globally, the plasma metabolome of VS-PWH was indistinguishable from PWoH, while Vi-PWH exhibited substantial metabolic disruption characterized by depletion of phosphatidylcholines, sphingomyelins, and hexosylceramides alongside triglyceride accumulation, dysregulation of the tryptophan-kynurenine and arginine-citrulline axes, and elevations in diacetylated polyamines and N4-acetylcytidine. Ordinal trend analysis identified subtle but consistent residual alterations in VS-PWH, particularly within phosphatidylcholine and sphingomyelin classes, that correlated with elevated TNFα and reduced CD4+ T cell counts and CD4/CD8 ratio. Together, these findings indicate that residual TNFα-associated inflammation and incomplete T cell recovery continue to shape the plasma metabolome in treated HIV, identifying candidate biomarkers for further mechanistic and clinical investigation.

## Introduction

Despite effective viral suppression on antiretroviral therapy (ART), people with HIV (PWH) continue to experience excess morbidity and mortality compared to people without HIV (PWoH), driven largely by an elevated burden of comorbidities including cardiovascular disease, certain cancers, and metabolic disorders such as diabetes^1,2^. Treatment options to mitigate these outcomes in virally suppressed (VS) PWH remain limited, in part because the underlying biological mechanisms are incompletely understood. Thus, there is an urgent unmet need for the characterization of biological factors driving poor health outcomes in virally suppressed PWH (VS-PWH).

Plasma metabolomics is a powerful tool for the identification of biomarkers relevant to disease pathogenesis and has been applied to HIV infection for over two decades, with early work identifying metabolites of tryptophan catabolism, such as kynurenine and quinolinic acid, as a hallmark of infection and correlate of disease progression^3,4^. Subsequent studies have gradually broadened this view to include lipids and other classes of small molecules, with a consensus that untreated HIV is associated with widespread metabolic disruption that partially resolves with ART^5,6^. Advances in high-dimensional targeted metabolomics now enable broader analyte coverage and the integration of metabolomic profiles with multiple immunologic and inflammatory measures in parallel, offering an opportunity to understand which metabolic features are restored by viral suppression and which persist, and how these disruptions relate to residual inflammation and T cell criteria (CD4+ T cell count, CD8+ T cell count, CD4/CD8 ratio, and nadir CD4+ T cell count).

In this study, we utilized high-dimensional targeted plasma metabolomics to quantify 254 small molecules and 750 lipids spanning 49 different classes along with 272 sum and ratio features and compared these between groups of viremic PWH (Vi-PWH), VS-PWH, and PWoH. We further evaluated a panel of plasma cytokines and investigated correlations between metabolomic features, cytokine concentrations, and T cell metrics with the goal of distinguishing metabolic features that are resolved by viral suppression from those that persist. Finally, we assessed relationships of residual metabolic disruption to specific markers of immune system health and inflammation.

## Results

### Differences in baseline characteristics

Baseline characteristics for study participants are shown in Table 1. Vi-PWH were significantly younger than both PWoH and VS-PWH (p < 0.001, respectively, Supplemental Figure 1A) and had been living with HIV for fewer years than VS-PWH (p < 0.001). There was an overall difference in gender between groups, with a significant pairwise difference between PWoH and VS-PWH (p = 0.0097, Supplemental Figure 1B), represented by more men in the virologically suppressed group. CD4+ T cell counts were significantly reduced in viremic individuals compared to both PWoH and VS-PWH (p < 0.001, respectively, Supplemental Figure 1C). There were no significant pairwise difference between groups for CD8+ T cell counts (Supplemental Figure 1D). The CD4/CD8 ratio was reduced in both VS-PWH and Vi-PWH compared to PWoH (p = 0.0012 and p < 0.001, respectively, Supplemental Figure 1E). Within VS-PWH, 100% of individuals were on an integrase strand transfer inhibitor (INSTI)-based regimen, with 85% being on a combination of bictegravir, tenofovir alafenamide, and emtricitabine.

**Table 1.** Baseline characteristics of enrolled participants.

|  | **PWoH**<br>(n = 17) | **VS-PWH**<br>(n = 20) | **Vi-PWH**<br>(n = 9) | **p-value** |
| --- | --- | --- | --- | --- |
| **Characteristic** |  |  |  |  |
| CD4<sup>+</sup> T cell count (cells/mm<sup>3</sup>) | 640 (170) | 488 (218) | 167 (190) | <0.001<sup>A</sup> |
| CD8<sup>+</sup> T cell count (cells/mm<sup>3</sup>) | 420 (174) | 623 (288) | 802 (718) | 0.034<sup>A</sup> |
| CD4/CD8 ratio | 1.88 (1.21) | 0.89 (0.39) | 0.20 (0.16) | <0.001<sup>A</sup> |
| Nadir CD4<sup>+</sup> T cell count (cells/mm<sup>3</sup>) | NA | 238 (201) | 106 (163) | 0.077<sup>B</sup> |
| Time living with HIV (years) | NA | 13.3 (5.7) | 3.0 (3.0) | <0.001<sup>B</sup> |
| Age (years) | 56 (14) | 52 (7) | 34 (7) | <0.001<sup>A</sup> |
| Sex at birth (n [%]) |  |  |  | 0.010<sup>C</sup> |
| Female | 9 (53%) | 2 (10%) | 1 (11%) |  |
| Male | 8 (47%) | 18 (90%) | 8 (89%) |  |
| Plasma HIV RNA (copies/mL) | NA | <LOD | 87,736 (144,809) |  |

Clinical characteristics of patients enrolled in the study. Continuous variables presented as mean (standard deviation).
<sup>A</sup>One-way analysis of variance (not assuming equal variances)
<sup>B</sup>Welch Two Sample t-test
<sup>C</sup>Fisher's exact test

### Antiretroviral therapy largely restores the plasma metabolome to HIV-negative phenotype

We first assessed global differences in plasma metabolome composition across groups by principal component analysis (PCA) and tested differences using pairwise PERMANOVAs (Figure 1). The plasma metabolome of virologically suppressed PWH did not differ significantly from PWoH (p = 0.295, Figure 1A), suggesting that effective viral suppression largely restores the plasma metabolome toward an HIV-negative phenotype. In contrast, viremic PWH exhibited significant metabolomic differences compared to both virologically suppressed PWH (p = 0.003) and PWoH (p = 0.003), indicating that active viremia is associated with substantial metabolic disruption. To further characterize the immunologic drivers of this variation, we examined the contribution of peripheral T cell criteria to plasma metabolome composition. Of the parameters tested, CD4+ T cell count explained the greatest proportion of variance and was the only significant predictor, outperforming CD8+ T cell count and the CD4/CD8 ratio (Figure 1B). Notably, the CD4/CD8 ratio explained an additional proportion of variance in the plasma metabolome after accounting for CD4+ and CD8+ T cell counts, although this contribution did not reach statistical significance. This suggests that the ratio captures aspects of immune composition not fully represented by either count alone. Together, these findings suggest that while ART effectively normalizes the plasma metabolome at a global level, there are significant metabolic disruptions in viremic PWH that appear more closely associated to CD4+ T cell depletion than to other T cell parameters.

**Figure 1.**
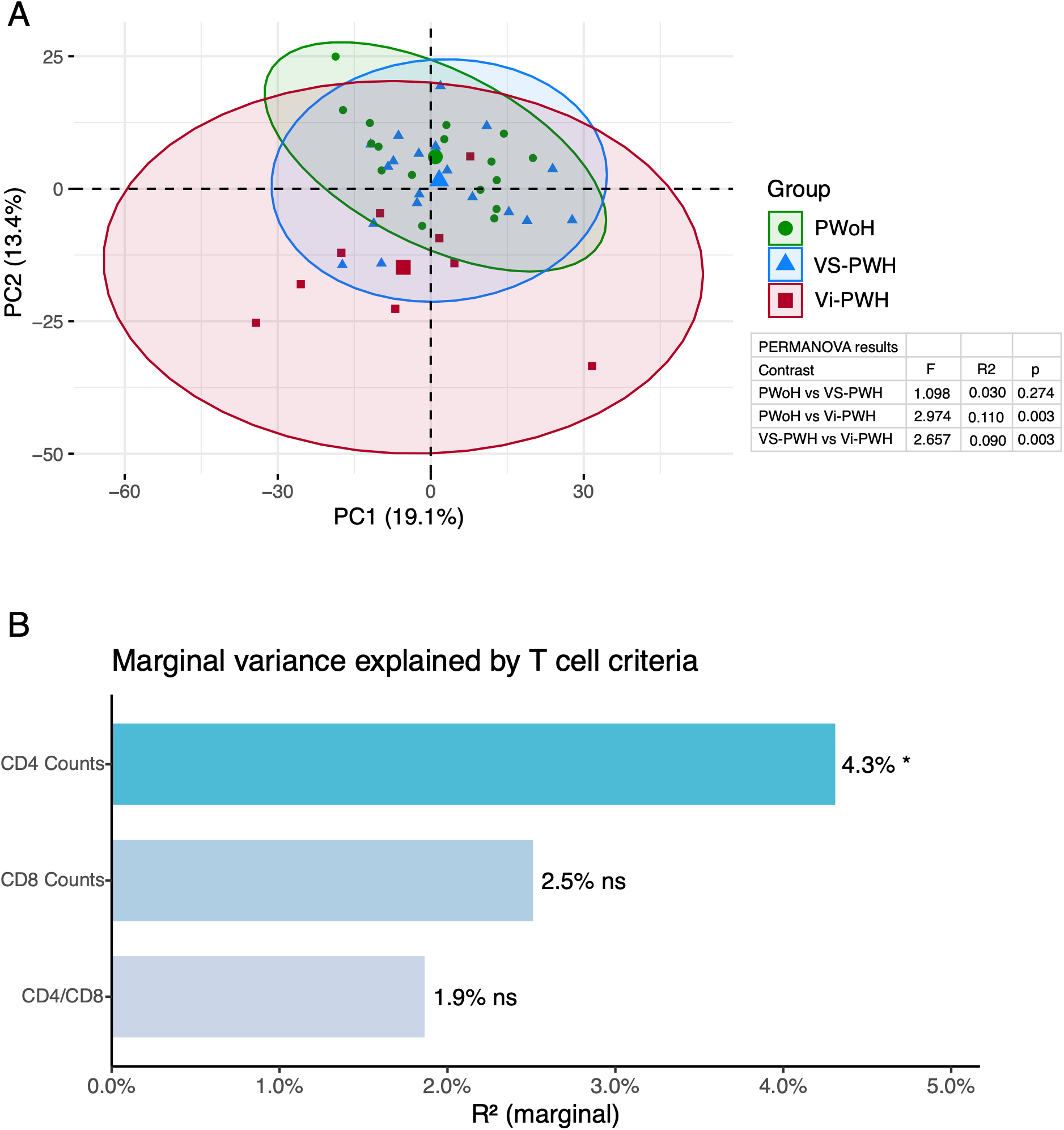
(A) Principal component analysis (PCA) of log2-transformed and scaled metabolite species. Each point represents an individual, with larger points indicating group centroids and ellipses representing 95% confidence intervals. Group differences were tested by PERMANOVA using a Euclidean distance matrix. (B) Proportion of variance in the Euclidean distance matrix independently explained by each T cell metric, assessed by marginal PERMANOVA.

### Differential abundance analysis reveals marked dyslipidemia during HIV viremia

Differential abundance testing of metabolite species (i.e. lipids and small molecules) and sum/ratio features across groups largely recapitulated the PCA results, with significant differences largely restricted to comparisons involving Vi-PWH (Figure 2). No metabolite species differed significantly between VS-PWH and PWoH after multiple comparison adjustment. In contrast, 150 species differed between Vi-PWH and PWoH, and 102 species differed between Vi-PWH and VS-PWH. Of these, 82 species were differentially abundant in both comparisons, while 68 were unique to the Vi-PWH vs. PWoH comparison and 20 were unique to the Vi-PWH vs. VS-PWH comparison (Figure 2B).

**Figure 2.**
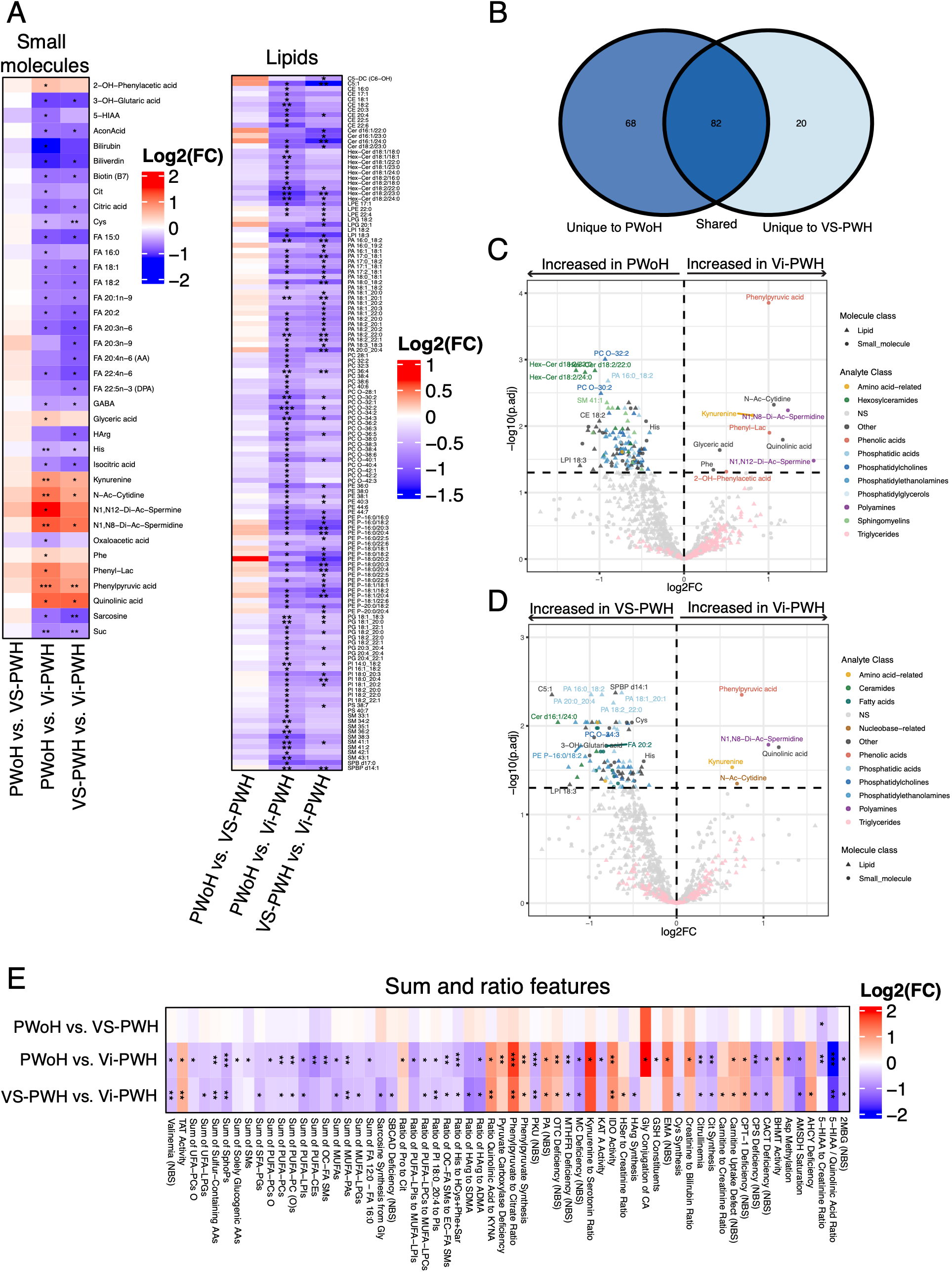
(A) Heatmap of metabolite species differentially abundant in at least one pairwise group comparison (FDR-adjusted p < 0.05), faceted by small molecule and lipid class. Columns represent pairwise contrasts; positive log2 fold-changes indicate higher abundance in the second listed group on the x-axis. (B) Overlap of differentially abundant species across pairwise comparisons. (C, D) Volcano plots of Vi-PWH versus PWoH (C) and Vi-PWH versus VS-PWH (D). (E) Heatmap of differentially abundant sum and ratio features, displayed as in (A). Asterisks denote the following FDR-adjusted p-values: * < 0.05, ** < 0.01, *** < 0.001.

The majority (78%) of differentially abundant species were lipids. Phosphatidylcholines (PCs), phosphatidylethanolamines (PEs), and phosphatidic acids (PAs) were broadly depleted in Vi-PWH relative to both PWoH and VS-PWH, while hexosylceramides (HexCers), phosphatidylglycerols (PGs), and sphingomyelins (SMs) were additionally elevated in PWoH compared to Vi-PWH (Figure 2C). Additional small molecules increased in both PWoH and VS-PWH compared to Vi-PWH included numerous fatty acids (FAs), especially polyunsaturated and long-chain unsaturated fatty acids within the linoleic acid pathway (i.e., linoleic acid, dihomo-γ-linolenic acid, arachidonic acid, and adrenic acid). Other notable small molecules depleted in Vi-PWH included bilirubin, biliverdin, biotin, citrulline, and gamma-Aminobutyric acid (GABA). In contrast, all metabolite species elevated in Vi-PWH were small molecules. Phenylpyruvic acid, quinolinic acid, kynurenine, N1, N8-diacetylspermidine, and N4-acetylcytidine were significantly elevated in Vi-PWH compared to both PWoH and VS-PWH (q < 0.05, Figures 2C–2D). Viremic PWH compared to PWoH additionally had elevated glyceric acid, phenyllactic acid, 2-hydroxyphenylacetic acid, phenylalanine, and N1, N12-diacetylspermine (q < 0.05, Figure 2C).

Sum and ratio features followed a similar pattern, with differences again restricted largely to Vi-PWH (Figure 2D). One exception was a decreased ratio of 5-hydroxyindoleacetic acid (5-HIAA) to creatinine in both Vi-PWH and VS-PWH relative to PWoH (q < 0.05), suggesting a reduction in serotonin catabolism that persists with viral suppression. Among Vi-PWH, indoleamine 2,3-dioxygenase (IDO) activity, tyrosine aminotransferase (TAT) activity, and phenylpyruvate synthesis were elevated relative to both comparison groups, while sums of monounsaturated fatty acids (MUFAs), MUFA-PAs, polyunsaturated fatty acid (PUFA) phosphatidylcholines, PUFA diacyl-phosphatidylcholines and sphingosine phosphates (SphoPs) were decreased (q < 0.05). Together, these findings indicate that HIV viremia is characterized by broad depletion of phospholipid and sphingolipid species alongside accumulation of polyamines, phenolic acids, and tryptophan catabolites, a metabolic profile largely restored with viral suppression.

### Metabolite set enrichment analysis reveals pattern of increasing hypertriglyceridemia with HIV severity

We next performed a metabolite set enrichment analysis (MSEA) to evaluate differences in KEGG pathway enrichment between groups (Figure 3). Comparing VS-PWH to PWoH, glycerolipid metabolism was significantly enriched while sphingolipid metabolism was depleted (q < 0.05, Figure 3A). The majority of metabolite species matching to the glycerolipid metabolism KEGG pathway in this analysis were triglycerides (TGs), indicating a broad pattern of TG accumulation in VS-PWH that was not captured by individual differential abundance testing. Sphingolipid pathway depletion was driven by HexCers, dihexosylceramides and trihexosylceramides, consistent with a broad reduction of glycosphingolipids in VS-PWH compared to PWoH.

**Figure 3.**
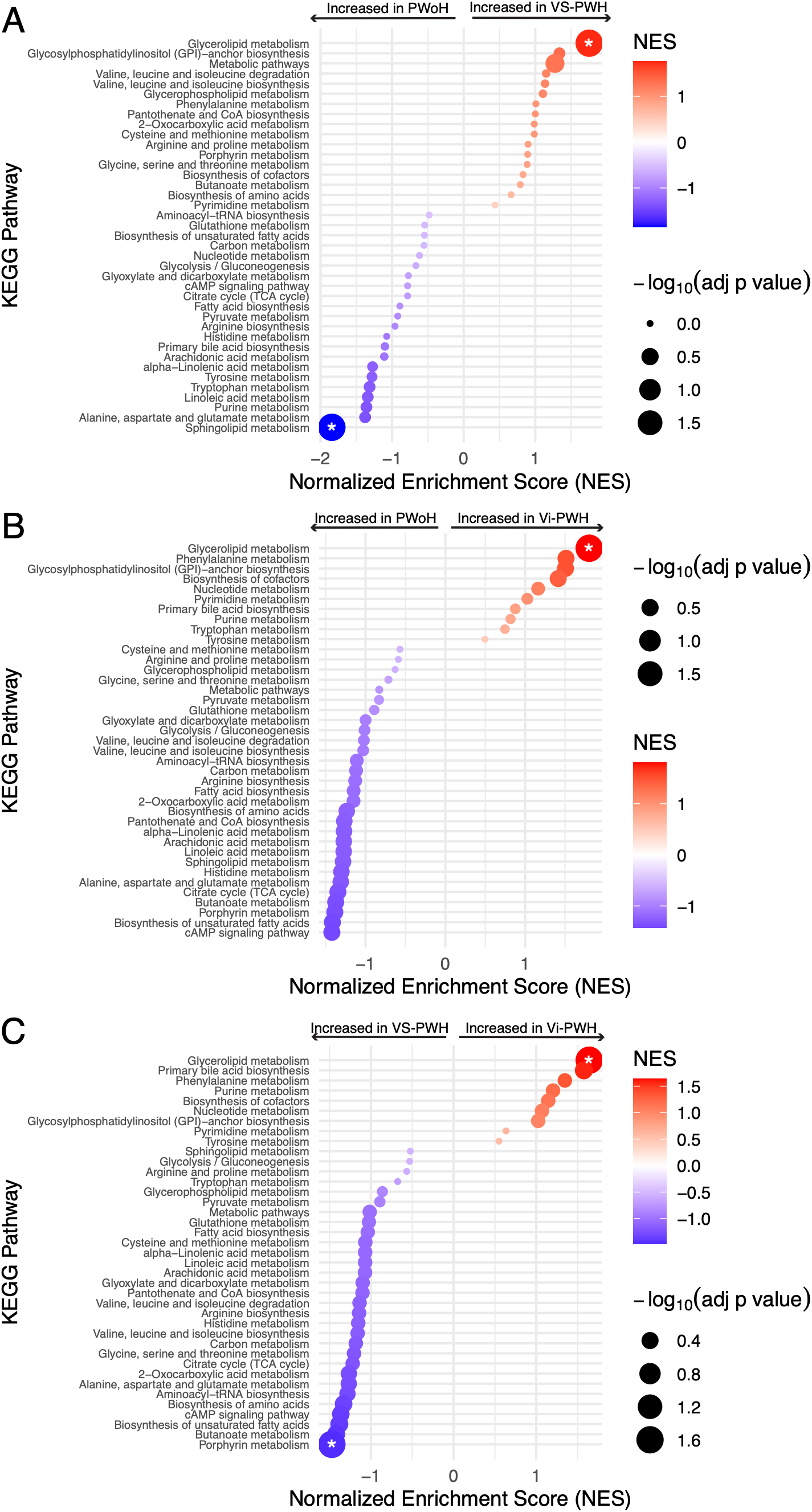
KEGG pathway enrichment by metabolite set enrichment analysis (MSEA) for VS-PWH versus PWoH (A), Vi-PWH versus PWoH (B), and Vi-PWH versus VS-PWH (C). Asterisks denote FDR-adjusted p-values < 0.05.

Glycerolipid metabolism was similarly enriched in Vi-PWH relative to both VS-PWH and PWoH (q < 0.05, Figures 3B–3C), with TGs again driving the leading edge, indicating that TG accumulation is a feature of HIV infection that does not fully resolve with viral suppression and intensifies during viremia. Porphyrin metabolism was additionally depleted in Vi-PWH compared to VS-PWH (q < 0.05, Figure 3C), with leading edge analysis indicating this was driven by reduced bilirubin and biliverdin.

### Elevated TNFɑ distinguishes viremic PWH

To better understand drivers of inflammation and metabolomic disruptions we evaluated cytokine concentrations between groups. Interestingly, despite the numerous differences observed in the plasma metabolome between Vi-PWH and VS-PWH, we did not observe any cytokine differences between these groups after adjusting for multiple comparisons (Figure 4A). The only significant difference we observed was increased TNFɑ in Vi-PWH compared to PWoH (q = 0.0067, Figure 4B). However, we additionally observed increased IL-10 and reduced IL-17A in Vi-PWH compared to PWoH that approached significance (q = 0.0582, respectively).

**Figure 4.**
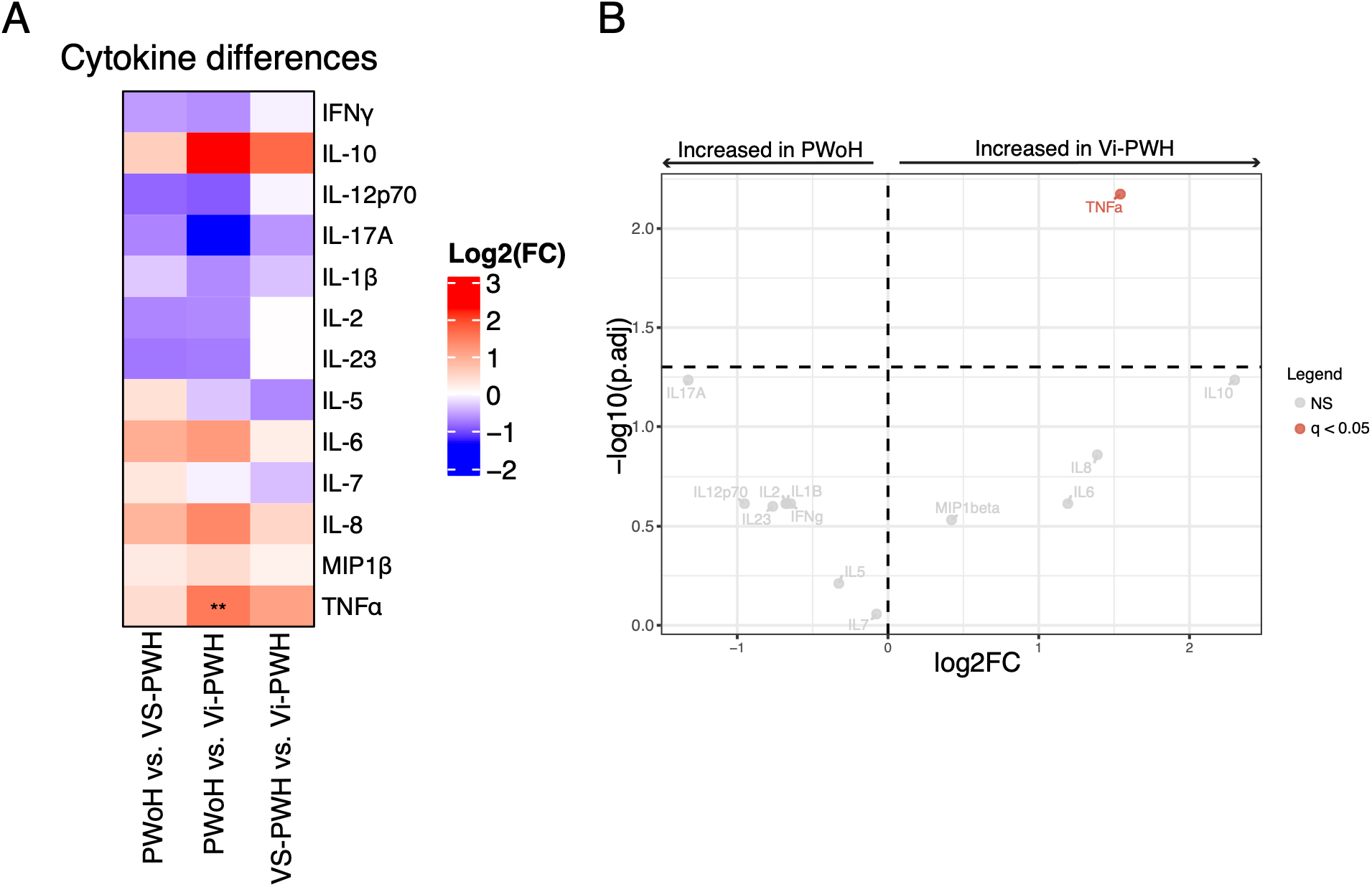
(A) Heatmap displaying differential abundance analysis of plasma cytokines between groups. Columns represent pairwise contrasts; positive log2 fold-changes indicate higher abundance in the second listed group on the x-axis. **p_FDR_ < 0.01. (B) Volcano plot of cytokine differences between PWoH and Vi-PWH, horizontal line represents cutoff for FDR-adjusted p-value < 0.05.

### Residual metabolic disruption persists in virally suppressed PWH

Given the absence of differentially abundant metabolites between VS-PWH and PWoH despite significant pathway-level enrichment differences, we next tested for metabolomic features that changed in an ordinal fashion across groups, from PWoH to VS-PWH to Vi-PWH. Such a pattern may indicate subtle metabolic changes that are not resolved during viral suppression. This analysis identified 43 metabolite species and 11 sum/ratio features that decreased monotonically with HIV disease severity, alongside only 2 metabolite species and 3 sum/ratio features that increased (q < 0.05, Figure 5A). Decreasing species were dominated by lipid classes including PCs, HexCers, SMs, and PGs, in addition to 5-HIAA and biliverdin. The two increasing species were both small molecules, phenylpyruvic acid and N4-acetylcytidine. PCA on only these trending metabolomic features shows increased separation between PWoH and VS-PWH, with significant PERMANOVA comparisons between all groups (q < 0.05, Figure 5B). The majority of separation between groups was along principal component 1, with the top loadings primarily being PCs and SMs (Figure 5C) and was positively correlated with CD4+ T cell counts (r = 0.56, q < 0.001), the CD4/CD8 ratio (r = 0.71, q < 0.001) and was negatively correlated with plasma TNFɑ (r = -0.49, q = 0.023, Figure 5D). Ordinal trend analysis further identified TNFα as the only cytokine that increased significantly with HIV disease severity (Figure 5E). Overall, these data suggest that viral suppression retains subtle metabolic disruptions compared to PWoH, primarily related to a loss of lipid classes (PCs and SMs), that appear to be driven by residual TNFɑ-associated inflammation.

**Figure 5.**
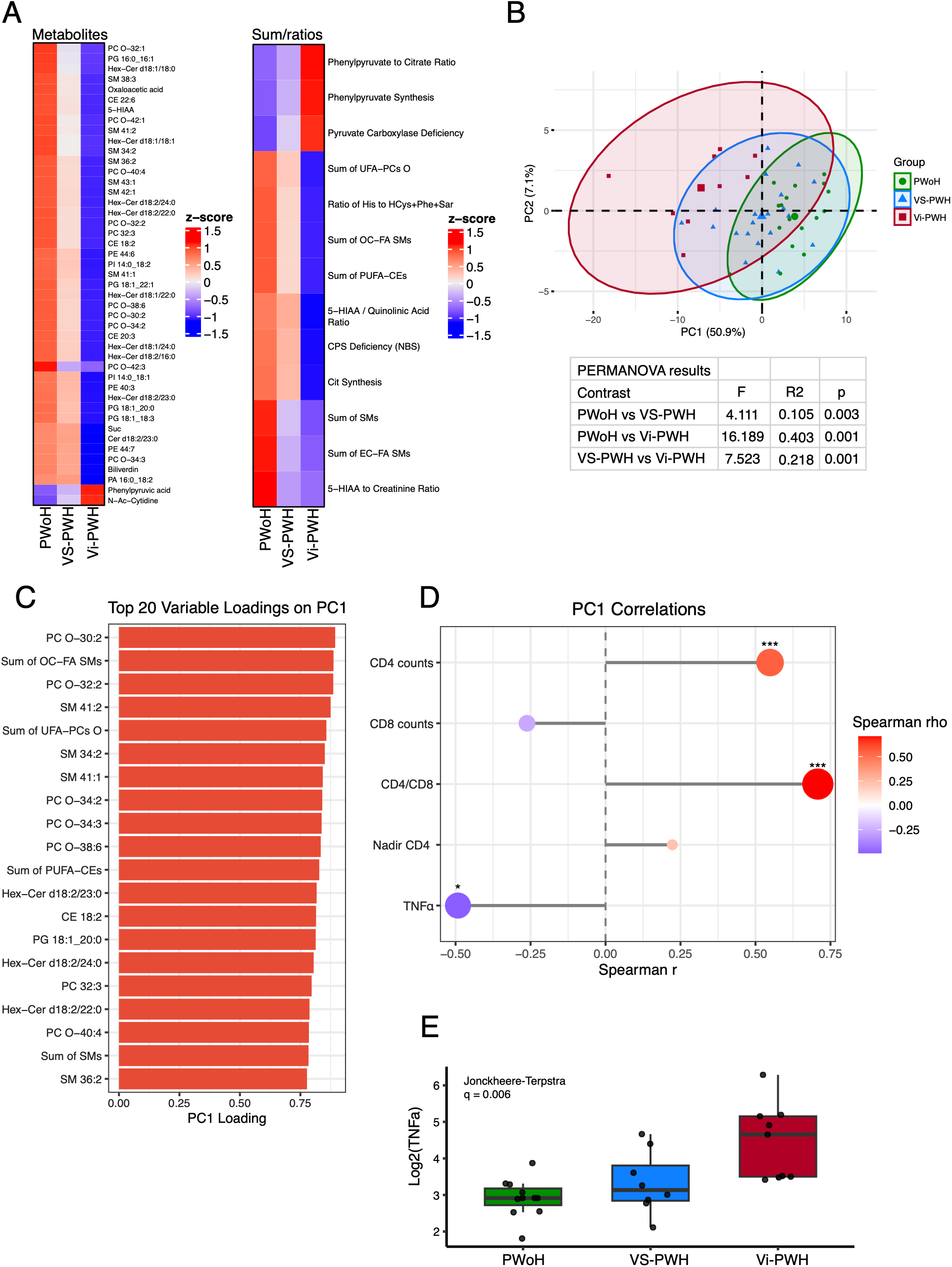
(A) Heatmap of features identified as ordinally increasing or decreasing across HIV severity groups by Jonckheere-Terpstra trend tests (FDR-adjusted p < 0.05), faceted by metabolite species and sum/ratio features. Cells represent group-average log2-transformed abundance, scaled by row. (B) PCA of ordinally trending features, with pairwise group differences tested by PERMANOVA on Euclidean distances. (C) Top 20 variable loadings on PC1 from (B). (D) Spearman correlations between PC1 and T cell criteria and TNFα. *q < 0.05, ***q < 0.001. (E) Plasma TNFα across HIV severity groups, with significance assessed by Jonckheere-Terpstra trend test.

### Metabolomic correlations with T cell criteria and TNFɑ

We next performed a correlation analysis between metabolomic features and T cell criteria. We identified 168 significant correlations between T cell criteria and metabolite species (q < 0.05, Supplemental Data 4, Figure 6A). To characterize the specificity of these associations, we compared the sets of metabolite nodes correlated with each T cell criterion (Figure 6B). Many of the correlated metabolites were shared between CD4+ T cell counts and the CD4/CD8 ratio (48 shared nodes), while the CD4/CD8 ratio was uniquely associated with an additional 50 metabolites/nodes. The nadir CD4 count was additionally uniquely correlated with 9 metabolites. Notably, CD8+ T cell counts only had a single correlated metabolite (SM 38:3) which was a shared node with the CD4/CD8 ratio. Most correlations with CD4+ T cell count and the CD4/CD8 ratio were positive and involved lipids, including PCs, PEs, SMs, and HexCers (Figure 6C), while nadir CD4 was generally positively correlated with fatty acids (e.g., docosapentaenoic acid, mead acid, pentadecanoic acid) and negatively correlated with polyamines (e.g., N1,N8-diacetylspermidine, N1, N12-diacetylspermine). Thus, better immune reconstitution is associated with a metabolic profile characterized by preserved lipid metabolism, increased fatty acids, and reduced metabolites linked to cellular stress, consistent with lower risk of ongoing immune dysfunction and non-AIDS comorbidities

**Figure 6.**
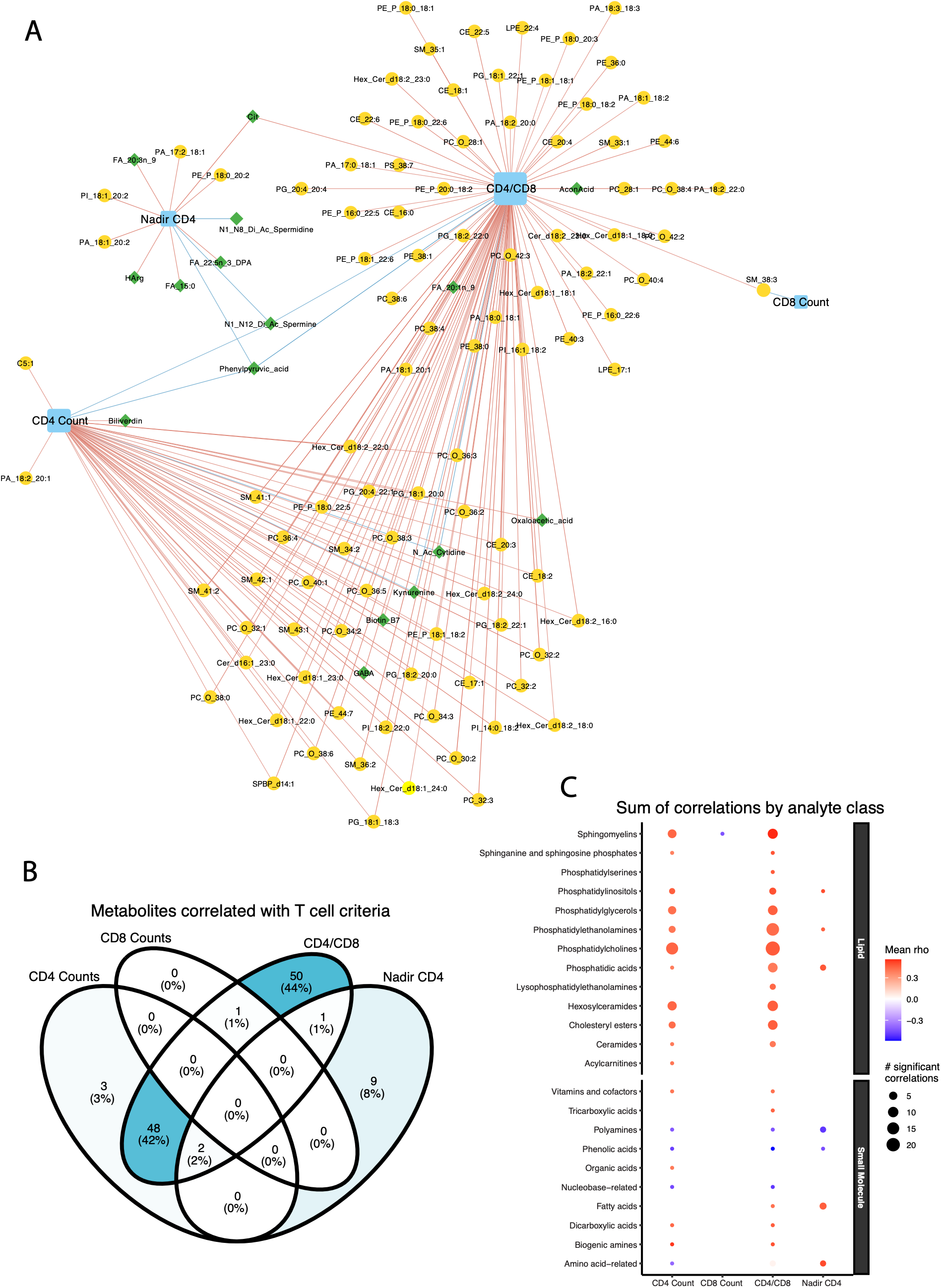
(A) Spearman correlation network between T cell criteria and metabolite species (FDR-adjusted p < 0.05). Edges are colored red (positive) or blue (negative); yellow nodes indicate lipid species and green nodes indicate small molecules. Node size reflects degree (number of correlations). (B) Overlap of metabolite species correlated with each T cell criteria. (C) Summary of correlations by analyte class, with each point representing an analyte class plotted by number of significant correlations and mean Spearman rho.

When looking at sum and ratio features, we identified 42 significant correlations with T cell criteria (q < 0.05, Supplemental Data 5, Figure 7A). Corroborating findings with metabolite species, we observed that the sum of lipid classes including PUFA cholesteryl esters (CEs), odd-chain fatty acid SMs, and unsaturated fatty acid acyl-alkyl-phosphatidylcholines were positively correlated with both the CD4 T cell count and CD4/CD8 ratio, whereas the CD4/CD8 ratio was uniquely positively correlated with the sum of several other lipid classes including SMs, PUFA phosphatidylcholines, PUFA diacyl-phosphatidylcholines, and SphoPs. The CD4/CD8 ratio was also negatively correlated with markers of IDO activity, carnitine uptake defect (ratio of C0, C2, C3, C16, C18, and C18:1 acylcarnitines to citrulline), and ornithine transcarbamylase (OTC) deficiency (ornithine-to-citrulline ratio). Further, the ratio of citrulline to ornithine and arginine was positively correlated with CD4+ T cell counts, the CD4/CD8 ratio, and nadir CD4, again demonstrating that the metabolome and immune restoration are associated.

**Figure 7.**
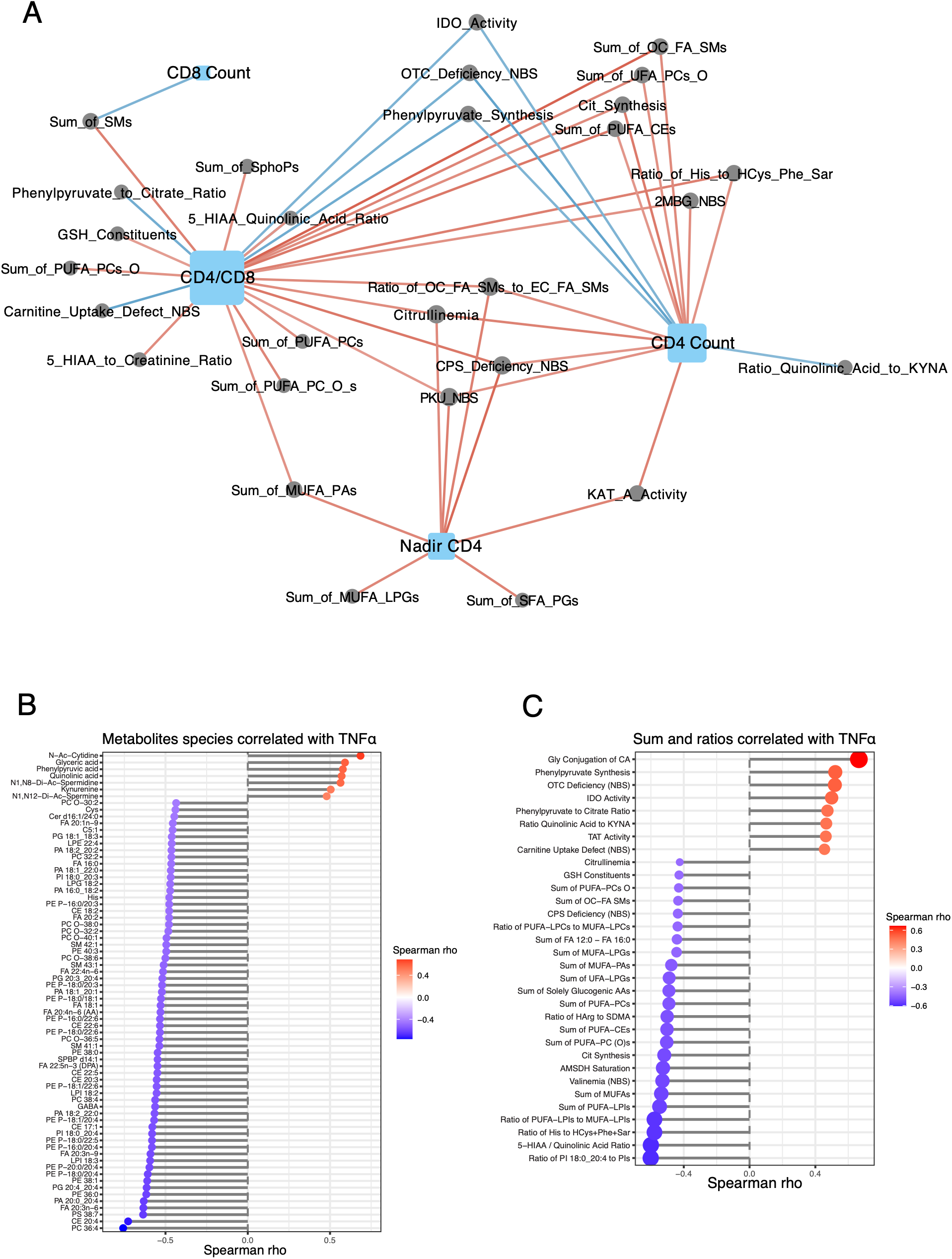
(A) Spearman correlation network between T cell criteria and metabolite sum/ratio features (FDR-adjusted p < 0.05). Edges are colored red (positive) or blue (negative). Node size reflects degree (number of correlations). (B) Summary of significant (FDR-adjusted p-value < 0.05) Spearman correlations between metabolite species and TNFα. (C) Summary of significant (FDR-adjusted p-value < 0.05) Spearman correlations between metabolite sum/ratio features and TNFα.

We next examined correlations between metabolomic features and TNFα, identifying 47 significant metabolite species and 32 sum/ratio features (q < 0.05, Figures 7B and 7C). TNFα was broadly negatively correlated with lipids spanning multiple classes (e.g., PEs, PCs, CEs), long-chain fatty acids in the linoleic acid pathway (e.g., dihomo-γ-linolenic acid, arachidonic acid, and adrenic acid), and GABA. In contrast, TNFα was positively correlated with tryptophan catabolites (kynurenine and quinolinic acid), diacetylated polyamines (N1,N8-diacetylspermidine and N1,N12-diacetylspermine), phenylpyruvic acid, glyceric acid, and N4-acetylcytidine. Sum and ratio features followed the same pattern: TNFα was negatively correlated with sums of PUFA cholesteryl esters and PUFA phosphatidylcholines, the 5-HIAA-to-quinolinic acid ratio, and citrulline synthesis (citrulline-to-ornithine ratio), and positively correlated with IDO activity, phenylpyruvate synthesis (phenylpyruvic acid-to-phenylalanine ratio), the phenylpyruvate-to-citrate ratio, and OTC deficiency. Together, these findings indicate that better T cell status and lower TNFα are linked through overlapping metabolic signatures, particularly enrichment of PUFA-containing lipids and intact citrulline synthesis, whereas elevated TNFα and worse T cell status share a shift toward phenylalanine catabolism, polyamine accumulation, and disrupted urea cycle and carnitine metabolism.

## Discussion

In this study, we used high dimensional targeted metabolomics to characterize the plasma metabolome of viremic PWH, virologically suppressed PWH, and PWoH, identifying distinct metabolic signatures associated with HIV disease state, T cell criteria, and systemic inflammation. At a global level, the plasma metabolome of VS-PWH was largely indistinguishable from that of PWoH, while Vi-PWH exhibited substantial metabolic disruption, consistent with the view that effective ART restores much, though not all, of the systemic perturbations caused by HIV^5,7^. CD4+ T cell count was the dominant immunologic correlate of metabolome composition, explaining more variance than either the CD4/CD8 ratio or CD8+ T cell count. Despite the apparent global normalization with viral suppression, ordinal trend analysis revealed that VS-PWH retain subtle but consistent metabolic alterations relative to PWoH, particularly across PC and SM classes, that are associated with increased TNFα and reduced CD4+ T cell counts and CD4/CD8 ratio. Together, these findings indicate that viral suppression alone is insufficient to fully the restore plasma metabolome and that residual TNFα-associated inflammation may continue to shape the metabolic landscape of treated HIV.

The most striking metabolic disruption in Vi-PWH was a duality of dyslipidemia, characterized by depletion of PCs, PEs, SMs, and HexCers contrasted with the accumulation of TGs. The simultaneous loss of structural phospholipids and accumulation of TGs is consistent with previous reports of dyslipidemia in untreated HIV^6^ and indicates remodeling of lipid metabolism during active viral replication. Several mechanisms likely contribute to the loss of phospholipid classes during viremia. First, HIV incorporates host PCs, SMs, and HexCers^8^ into its envelope particle during budding, which may consume the circulating pool of these lipids over time. Second, elevated TNFα, driven by microbial translocation, co-infections, or active HIV replication, increases phospholipase A2 (PLA2) activity which hydrolyzes glycerophospholipids (e.g., PCs, PEs), producing arachidonic acid and lysophospholipids that can further amplify inflammatory signaling through the production of eicosanoids^9,10^. In parallel, accumulation of TGs likely reflects both reduced clearance and increased synthesis driven by inflammatory processes. TNFα increases free fatty acid (FFA) production in the adipose tissue and liver, which serve as precursors for TGs, while also suppressing lipoprotein lipase (LPL) activity which results in decreased clearance of TG-rich very low-density lipoproteins (VLDLs)^11^. Importantly, MSEA revealed that the KEGG glycerolipid pathway was most enriched in Vi-PWH, followed by VS-PWH compared to PWoH, indicating that TG accumulation is a feature of HIV infection that intensifies during viremia but does not fully resolve with ART. This pattern aligns with the well-documented elevated cardiovascular and metabolic disease risk (e.g., diabetes) in PWH on long-term ART^2^.

A second notable feature of Vi-PWH was disruption of the tryptophan-kynurenine pathway. Elevations in kynurenine and quinolinic acid, together with increased IDO activity (ratio of kynurenine to tryptophan), are well-established hallmarks of chronic HIV infection and have been linked to T cell dysfunction, disease progression, and is also associated with microbial dysbiosis in HIV infection^12–14^. This was accompanied by decreased flux through the trypophan-serotonin pathway, as evidenced by reduced 5-HIAA concentrations in Vi-PWH, which also significantly trended down with HIV severity suggesting that this disruption is not fully resolved on ART. We also observed disruption of the urea cycle and arginine-citrulline axis. Citrulline was reduced in Vi-PWH, and ratios reflecting impaired citrulline synthesis (citrulline-to-ornithine) and OTC dysfunction (ornithine-to-citrulline) were negatively associated with the CD4/CD8 ratio. Citrulline synthesis additionally trended down with HIV severity, indicating this axis continues to be disrupted on ART. Plasma citrulline is produced almost exclusively by small intestine enterocytes and serves as a marker of enterocyte mass and function, making these findings consistent with the gut epithelial damage and microbial translocation that characterize chronic HIV^15^. The negative correlation between TNFα and citrulline synthesis, as well as positive correlations between citrulline, nadir CD4, and the CD4/CD8 ratio further supports that systemic inflammation, immune activation, and damage to the gut mucosa are interrelated.

An increase of diacetylated polyamine species represented another major feature of metabolic disruption in Vi-PWH. N1,N8-diacetylspermidine and N1,N12-diacetylspermine were among the most significantly elevated metabolites in Vi-PWH and were both negatively correlated with nadir CD4 count, indicating that the accumulation of these polyamines is associated with historical immune damage. N1,N12-diacetylspermine was further positively correlated with TNFα, and negatively correlated with both CD4+ T cell counts and the CD4/CD8 ratio, indicating its abundance may also be related to inflammation and immune activation. Polyamines, including putrescine, spermine, and spermidine have essential functions for many processes including cellular proliferation, protein synthesis, and cell death^16^. Given these essential roles, intracellular polyamine concentrations are tightly regulated, in part through catabolism by spermidine/spermine N1-acetyltransferase (SSAT/SAT1), the rate-limiting enzyme that acetylates spermidine and spermine to produce acetylated intermediates. These intermediates are then oxidized by polyamine oxidase (PAOX) to regenerate lower polyamines, producing reactive oxygen species (ROS) in the process^17^. Diacetylated polyamines in particular accumulate from increased SSAT activity^18^. SSAT can be induced by several factors, including type I interferon responses^19^ and the HIV Tat protein, with associations between increased SSAT activity and HIV-associated neurocognitive decline^20^. We did not observe differences in other plasma polyamines (e.g., spermidine and putrescine), suggesting that the elevated diacetylated polyamines reflect increased cellular export driven by viremia-mediated SSAT activity. However, we did not measure intracellular polyamines and cannot directly assess intracellular polyamine metabolism. Notably, both N1,N8-diacetylspermidine and N1,N12-diacetylspermine have been described as cancer biomarkers, raising the possibility that they may be informative in PWH given the elevated HIV-associated cancer risk^21–23^.

Another acetylated metabolite elevated in Vi-PWH was N4-acetylcytidine, a modified nucleoside generated when N-acetyltransferase 10 (NAT10) acetylates cytidine residues on RNA^24^. N4-acetylcytidine plays an important role in translation but has also been linked to a variety of human diseases, most commonly cancer^24,25^. Interestingly, N4-acetylcytidine has more recently been recognized for its role in viral infections through stabilization of viral RNA. The HIV virus for instance, was recently shown to utilize NAT10 activity to modify viral RNA, leading to increased viral RNA stability and replication^26^. In plasma, N4-acetylcytidine concentration likely reflects whole-body RNA turnover, as has been described for urinary measurements^27^. Elevated circulating N4-acetylcytidine in Vi-PWH may therefore be derived from contributions of both viral and host cell RNA turnover during active replication. Consistent with this, we observed that N4-acetylcytidine was strongly positively correlated with TNFα, negatively correlated with CD4+ T cell counts and the CD4/CD8 ratio, and was one of the few metabolite species that trended upward with HIV severity, pointing to an inflammation-associated effect that is not fully resolved on ART.

Our correlation analysis revealed distinct relationships between the plasma metabolome and individual T cell metrics. The CD4/CD8 ratio was the most highly interconnected, correlating with 126 metabolomic features, compared to 68 for CD4+ T cell counts, 20 for nadir CD4, and just 2 for CD8+ T cell counts. Notably, only 4 of the CD4+ T cell count correlations were unique to that metric, the remainder were shared with the CD4/CD8 ratio or nadir CD4. These findings underscore both the complementary information captured by each T cell metric and the value of the CD4/CD8 ratio as a clinical indicator. These data further point to the importance of preserving T cell health to limit metabolomic disruption, particularly through early ART initiation and minimization of immune activation. However, it is unclear which is cause and effect here and further studies to investigate this are warranted. Associations with TNFα primarily highlight the negative influence of TNFα on lipid classes as well as long-chain fatty acids, particularly those in the linolenic acid pathway. These findings suggest that interventions targeting the drivers of increased TNFα, possibly including microbial translocation, co-infections, and HIV reservoir activity, may help mitigate the persistent metabolic disruptions observed in treated HIV^28–31^.

This study must be interpreted in the context of its limitations. First, the observational and cross-sectional design precludes causal inference. Next, the relatively modest sample size, combined with the high dimensionality of the dataset, limits power to detect subtle differences within the VS-PWH group. We therefore relied on pathway enrichment and identification of features that changed ordinally with HIV severity to characterize residual metabolic disruption in VS-PWH. The effect sizes reported here (Supplemental Data 1-3) may help inform the design of future, better-powered and more targeted studies of metabolomic differences in treated HIV. Our analyses were also not adjusted for covariates such as age, sex, diet, BMI, medication use, or comorbidities, owing both to incomplete metadata and to confounding between groups for variables such as age and sex. Our results should therefore be regarded as exploratory and hypothesis-generating. We are also unable to distinguish ART-related from HIV-related contributions to the metabolic differences observed in VS-PWH. The fact that metabolites trending with HIV severity in VS-PWH more closely resemble those of Vi-PWH, who are not currently virally suppressed on ART, suggests a predominantly HIV-associated effect, but longitudinal studies including ART initiation will be needed to confirm this. Further, a larger cohort to disentangle VS-PWH that are at high and low risk for serious non-AIDS events is warranted. Finally, integration of other omic-layers, such as the gut microbiome, co-infection status, and HIV reservoir measurements would allow for a deeper understanding of the drivers of residual inflammation. Despite these limitations, our findings provide a detailed view of the plasma metabolome across a spectrum of HIV severity and identify persistent metabolic disruptions in virally suppressed individuals that warrant further mechanistic and clinical investigation.

In summary, this high-dimensional targeted metabolomics study of viremic PWH, virally suppressed PWH, and PWoH demonstrates that effective ART largely restores the plasma metabolome to an HIV-negative phenotype except for a subtle signature of metabolic disruption related to ongoing inflammation and incomplete immune recovery. Viremia was associated with broad metabolic disruption, including a dyslipidemic pattern of phospholipid and sphingolipid depletion alongside triglyceride accumulation, dysregulation of the tryptophan-kynurenine and arginine-citrulline axes, and elevations in diacetylated polyamines and N4-acetylcytidine. Although viral suppression normalized the global metabolome, subtle but consistent alterations persisted in VS-PWH, particularly within glycerophospholipid and sphingomyelin classes, which correlated with TNFα and T cell criteria. Together, this work indicates that residual TNFα-associated inflammation and incomplete T cell recovery continue to shape the plasma metabolome in treated HIV. Here, by defining the metabolic consequences of residual inflammation and incomplete immune recovery, these findings establish a foundation for biomarker-guided risk prediction and the development of targeted interventions to improve long-term outcomes in people with HIV.

## Methods

### Participant enrollment

Participants were enrolled at the Center for Research in Infectious Disease (CIENI) of the National Institute of Respiratory Diseases (INER) in Mexico City. All PWH were confirmed HIV-positive. Viremic individuals had all been previously diagnosed with HIV prior to enrollment but were non-adherent to ART upon screening.

### Study approval

All participants gave written informed consent using IRB-approved forms. The INER Ethics and Research Committees approved the study (C71-18) in Mexico.

### Sample collection

Blood samples were collected in EDTA tubes and plasma was isolated by density gradient centrifugation. Plasma samples were aliquoted and stored at -80°C until used for our downstream analyses.

### Plasma metabolomics

Plasma metabolomic profiling was performed by biocrates using the MxP Quant 1000 kit. Lipid species were quantified by flow injection analysis coupled to tandem mass spectrometry (FIA-MS/MS) on a 5500 QTRAP platform (AB Sciex, Darmstadt, Germany) equipped with an electrospray ionization source, while small molecule analytes were quantified by liquid chromatography-tandem mass spectrometry (LC-MS/MS) on a 5500+ Triple Quad instrument (AB Sciex, Darmstadt, Germany). Prior to statistical analysis, metabolites detected in fewer than 50% of samples were excluded, and values falling below the lower limit of detection were imputed as half the minimum detectable concentration. A complete list of quantified metabolites and formulas for sum and ratio features are provided in Supplemental Data 6-7.

### Plasma protein measurements

Cytokine concentrations in plasma were measured using a 13-plex (IL-1β, IL-2, IL-5, IL-6, IL-7, IL-8, IL-10, IL-12p70, IL-17A, IL-23, TNFα, IFNγ, and MIP-1β) Luminex panel (Cat. HSTCMAG-28SK, MILLIPLEX). Values below the limit of detection were imputed as half the minimum detectable concentration.

### Statistics

All statistical analysis was performed in R (version 4.4.1). Metabolomic features (including sum/ratios) were log_2_ transformed prior to all downstream analysis. Differences in baseline characteristics across groups were assessed using one-way ANOVA for continuous variables and Fisher’s exact test for categorical variables, with pairwise comparisons between groups performed using Tukey’s honest significant difference test and Fisher’s exact test, respectively. Differences in overall plasma metabolome composition were tested by pairwise PERMANOVA using the Euclidean distance matrix. Differential abundance testing was performed using the limma R package, requiring an FDR-adjusted p-value ≤ 0.05 for significance and adjusting separately between each contrast. MSEA was performed using HMDB-matched identifiers for all available metabolites, supplied as a pre-ranked list ordered by the product of the nominal p-value and log_2_ fold change. Enrichment analysis was carried out using the multiGSEA and fgsea R packages against KEGG pathways, requiring a minimum set size of 5 and statistical significance defined as an FDR-adjusted p-value ≤ 0.05. Trend analysis was performed by Jonckheere-Terpstra tests, assessing for features that ordinally increased or decreased from PWoH, virally suppressed PWH, and viremic PWH with FDR p-value adjustment performed separately for each direction. Spearman correlations were performed between metabolomic features, T cell criteria, and TNFα concentrations. To limit comparisons, only features that were determined as differentially abundant from the limma analysis were included.

## Data availability

Deidentified data and analysis code available from corresponding author upon reasonable request.

## Contributions

Conceptualization: NRK, TWS, GS, CMB

Supervision: NRK, TWS, GS, SAR, JA, CMB, MLG

Funding acquisition: NRK, TWS, GS

Investigation: CMB, JA, KE, GW, CG, TS, SAR, FTR, MSN, LCR, KKOC, OB, MLG, CH

Formal analysis: CMB

Visualization: CMB

Writing – original draft: CMB

Writing – review & editing: NRK, ES, RTC, NF, KE, SAR, TWS

## Acknowledgements

We are deeply grateful to the participants of this study. We would like to thank biocrates for generating the metabolomic data and the UMN Preclinical Research Center for generating the cytokine data.

## Abbreviations

5-HIAA: 5-hydroxyindoleacetic acid
ART: Antiretroviral therapy
BMI: Body mass index
CIENI: Center for Research in Infectious Disease
CE: Cholesteryl ester
FDR: False discovery rate
FA: Fatty acid
FIA-MS/MS: Flow injection analysis coupled to tandem mass spectrometry
FFA: Free fatty acid
GABA: Gamma-aminobutyric acid
HexCer: Hexosylceramide
HIV: Human immunodeficiency virus
HMDB: Human Metabolome Database
IDO: Indoleamine 2,3-dioxygenase
INSTI: Integrase strand transfer inhibitor
IL: Interleukin
KEGG: Kyoto Encyclopedia of Genes and Genomes
LC-MS/MS: Liquid chromatography-tandem mass spectrometry
LPL: Lipoprotein lipase
MSEA: Metabolite set enrichment analysis
MUFA: Monounsaturated fatty acid
NAT10: N-acetyltransferase 10
INER: National Institute of Respiratory Diseases
OTC: Ornithine transcarbamylase
PWH: People with HIV
PWoH: People without HIV
PERMANOVA: Permutational multivariate analysis of variance
PA: Phosphatidic acid
PC: Phosphatidylcholine
PE: Phosphatidylethanolamine
PG: Phosphatidylglycerol
PLA2: Phospholipase A2
PAOX: Polyamine oxidase
PUFA: Polyunsaturated fatty acid
PCA: Principal component analysis
ROS: Reactive oxygen species
RNA: Ribonucleic acid
SM: Sphingomyelin
SphoP: Sphingosine phosphate
SSAT/SAT1: Spermidine/spermine N1-acetyltransferase
TG: Triglyceride
TNFα: Tumor necrosis factor alpha
TAT: Tyrosine aminotransferase
VLDL: Very low-density lipoprotein
Vi-PWH: Viremic people with HIV
VS: Virologically suppressed

---

**Supplemental Figure 1.**
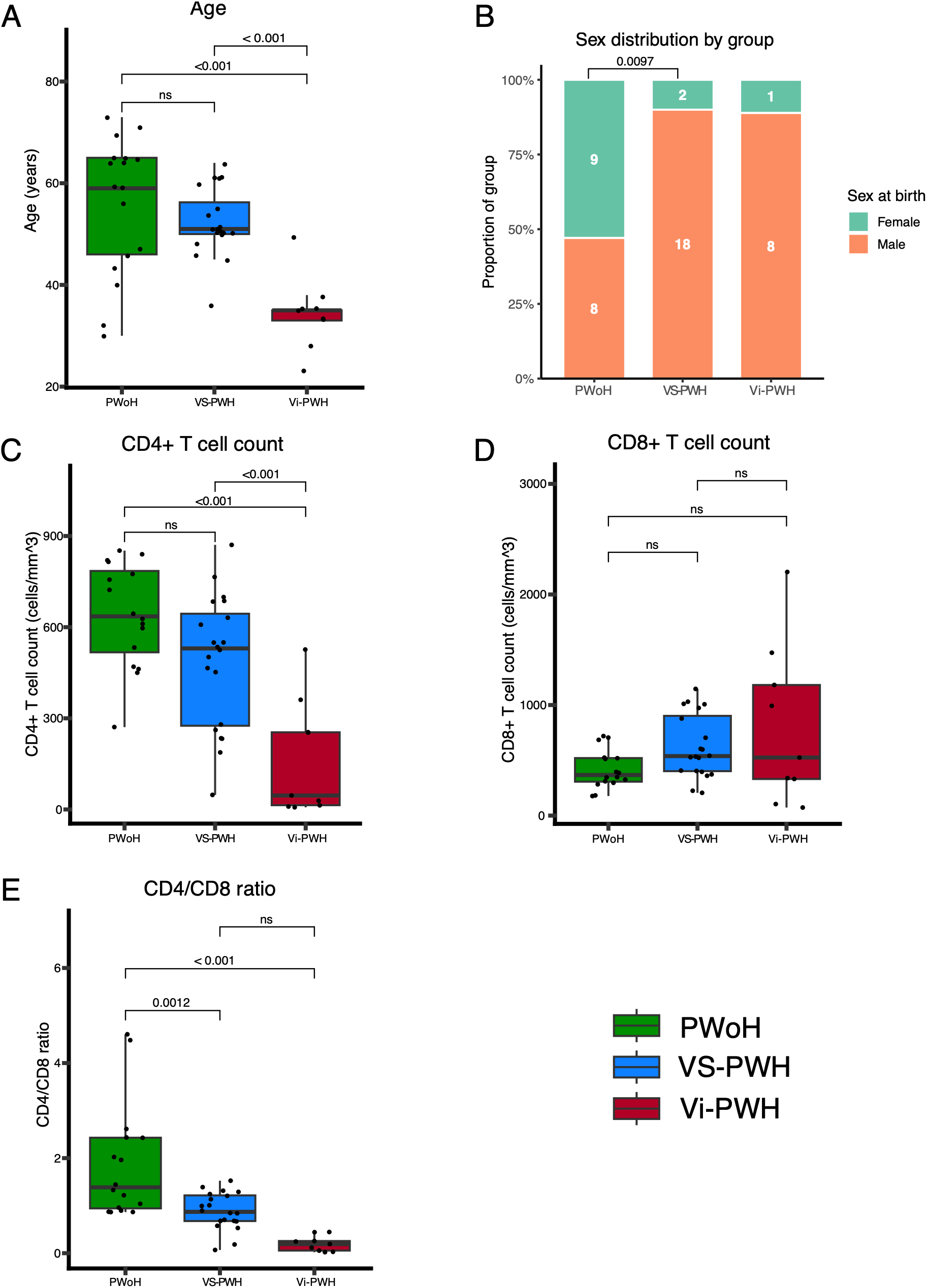
Differences between groups in age (A), sex at birth (B), CD4+ T cell count (C), CD8+ T cell count (D), and CD4/CD8 ratio (E). Pairwise comparisons for continuous variables were performed using Tukey’s honest significant difference (HSD) test; sex was compared using pairwise Fisher’s exact tests. Boxplots represent the median and 25^th^-75th percentiles, with whiskers extending to the largest and smallest values within 1.5 times the interquartile range.

## References

1. Trickey, A. et al. Life expectancy after 2015 of adults with HIV on long-term antiretroviral therapy in Europe and North America: a collaborative analysis of cohort studies. Lancet HIV 10, e295–e307 (2023).

2. Lerner, A. M., Eisinger, R. W. & Fauci, A. S. Comorbidities in Persons With HIV: The Lingering Challenge. JAMA 323, 19–20 (2020).

3. Heyes, M. P. et al. Quinolinic acid in cerebrospinal fluid and serum in HIV-1 infection: relationship to clinical and neurological status. Ann. Neurol. 29, 202–209 (1991).

4. Huengsberg, M. et al. Serum kynurenine-to-tryptophan ratio increases with progressive disease in HIV-infected patients. Clin. Chem. 44, 858–862 (1998).

5. Virseda-Berdices, A. et al. Plasma metabolomic profile is near-normal in people with HIV on long-term suppressive antiretroviral therapy. Front. Cell. Infect. Microbiol. 14, (2024).

6. Bowman, E. R. et al. Plasma lipidome abnormalities in people with HIV initiating antiretroviral therapy. Transl. Med. Commun. 5, 26 (2020).

7. Hileman, C. O. & Funderburg, N. T. Inflammation, Immune Activation, and Antiretroviral Therapy in HIV. Curr. HIV/AIDS Rep. 14, 93–100 (2017).

8. Lorizate, M. et al. Comparative lipidomics analysis of HIV-1 particles and their producer cell membrane in different cell lines. Cell. Microbiol. 15, 292–304 (2013).

9. Knuplez, E. & Marsche, G. An Updated Review of Pro- and Anti-Inflammatory Properties of Plasma Lysophosphatidylcholines in the Vascular System. Int. J. Mol. Sci. 21, 4501 (2020).

10. Wang, B. et al. Metabolism pathways of arachidonic acids: mechanisms and potential therapeutic targets. Signal Transduct. Target. Ther. 6, 94 (2021).

11. Popa, C., Netea, M. G., van Riel, P. L. C. M., van der Meer, J. W. M. & Stalenhoef, A. F. H. The role of TNF-α in chronic inflammatory conditions, intermediary metabolism, and cardiovascular risk. J. Lipid Res. 48, 751–762 (2007).

12. Zhang, J. et al. Amino acid metabolism dysregulation associated with inflammation and insulin resistance in HIV-infected individuals with metabolic disorders. Amino Acids 55, 1545–1555 (2023).

13. Favre, D. et al. Tryptophan catabolism by indoleamine 2,3-dioxygenase 1 alters the balance of TH17 to regulatory T cells in HIV disease. Sci Transl Med 2, 32ra36 (2010).

14. Vujkovic-Cvijin, I. et al. Dysbiosis of the gut microbiota is associated with HIV disease progression and tryptophan catabolism. Sci Transl Med 5, 193ra91 (2013).

15. Bahri, S. et al. Citrulline: From metabolism to therapeutic use. Nutrition 29, 479–484 (2013).

16. Rossi, M. N. & Cervelli, M. Polyamine Metabolism and Functions: Key Roles in Cellular Health and Disease. Biomolecules 14, 1570 (2024).

17. Casero, R. A., Jr & Pegg, A. E. Polyamine catabolism and disease. Biochem. J. 421, 323–338 (2009).

18. Kramer, D. L. et al. Polyamine Acetylation Modulates Polyamine Metabolic Flux, a Prelude to Broader Metabolic Consequences*. J. Biol. Chem. 283, 4241–4251 (2008).

19. Huang, M., Zhang, W., Chen, H. & Zeng, J. Targeting Polyamine Metabolism for Control of Human Viral Diseases. Infect. Drug Resist. 13, 4335–4346 (2020).

20. Merali, S. et al. Polyamines: Predictive Biomarker for HIV-Associated Neurocognitive Disorders. J. AIDS Clin. Res. 5, 1000312 (2014).

21. Hiramatsu, K. et al. Diagnostic and prognostic usefulness of N1,N8-diacetylspermidine and N1,N12-diacetylspermine in urine as novel markers of malignancy. J. Cancer Res. Clin. Oncol. 123, 539–545 (1997).

22. Kawakita, M. & Hiramatsu, K. Diacetylated derivatives of spermine and spermidine as novel promising tumor markers. J. Biochem. (Tokyo) 139, 315–322 (2006).

23. Basting, C. M. et al. Multi-omics links microbial dysbiosis, systemic inflammation and metabolomic disruptions to SNAE risk in treated HIV. 2026.04.09.717347 Preprint at 10.64898/2026.04.09.717347 (2026).

24. Jin, G., Xu, M., Zou, M. & Duan, S. The Processing, Gene Regulation, Biological Functions, and Clinical Relevance of N4-Acetylcytidine on RNA: A Systematic Review. Mol. Ther. Nucleic Acids 20, 13–24 (2020).

25. Zhang, S. et al. Recent advances in the potential role of RNA N4-acetylcytidine in cancer progression. Cell Commun. Signal. 22, 49 (2024).

26. Tsai, K. et al. Acetylation of Cytidine Residues Boosts HIV-1 Gene Expression by Increasing Viral RNA Stability. Cell Host Microbe 28, 306–312.e6 (2020).

27. Seidel, A., Brunner, S., Seidel, P., Fritz, G. I. & Herbarth, O. Modified nucleosides: an accurate tumour marker for clinical diagnosis of cancer, early detection and therapy control. Br. J. Cancer 94, 1726–1733 (2006).

28. Basting, C. M. & Klatt, N. R. Dissecting the Impact of the Gut Microbiome on HIV Reservoir Dynamics. J. Infect. Dis. 233, 622–624 (2026).

29. Swanson, E. C., Basting, C. M. & Klatt, N. R. The role of pharmacomicrobiomics in HIV prevention, treatment, and women’s health. Microbiome 12, 254 (2024).

30. Crawford, P. A. et al. Ongoing lymphoid HIV production drives pyroptosis and GLP-1 counter-regulation in ART-suppressed infection. 2026.01.09.698696 Preprint at 10.64898/2026.01.09.698696 (2026).

31. Chakrawarti, A. et al. Pre-treatment Microbiome Diversity and Function is associated with Expansion of Cytotoxic and Regulatory Immune Populations after N-803 treatment in People with HIV. 2025.10.01.679827 Preprint at 10.1101/2025.10.01.679827 (2025).

